# Thermal pre-treatment of algal symbiont species differentially affects coral development

**DOI:** 10.64898/2026.07.24.739192

**Authors:** Maria Ruggeri, Samuel Bedgood, Jun B. Cai, Jennifer Qian, Federica Montesanto, Mark McCauley, Grayce E. Dyer, Ayobami Oluokun, Mojibola Fowowe, Odunayo Oluokun, Yehia Mechref, Saki Harii, Sandra Loesgen, Virginia M. Weis

## Abstract

The nutritional relationship between coral hosts and intracellular algal symbionts is foundational to coral reef ecosystems. Although these symbioses are obligate for coral, many are re-established anew each host generation. Furthermore, climate change destabilizes the symbiosis, and the cellular mechanisms underlying symbiont colonization and host development, and how they are affected by thermal stress are poorly understood. Here, we investigated the effect of algal species and thermal treatments on symbiont and host cell proliferation by offering *Acropora tenuis* larvae one of four algal species pre-exposed to elevated or ambient temperature. We also characterized the cell-surface glycome of each species-temperature combination to understand its role in symbiont recognition and proliferation. We found that thermal pre-treatment negatively affected algal photosynthetic efficiency and initial symbiont density but did not affect symbiont colonization. In contrast, host cell proliferation was affected in a species-specific manner. Thermal pre-treatment of *Breviolum minutum* and *Durusdinium trenchii* negatively affected host cell proliferation, whereas thermal treatment of *Symbiodinium microadriaticum* did not affect development. Further, uptake of thermally pre-treated *D. trenchii* decreased host cell proliferation below that of larvae without symbionts, indicating a cost to host development despite this species’ high thermal tolerance. Algal glycans varied across species but not thermal pre-treatments, suggesting reductions in density of thermally pre-treated algae may be due to changes in physiology rather than altered surface chemistry. Further, variation in glycan abundance across species did not track differences in colonization rate or symbiont density, hinting towards a smaller role of glycans in host-symbiont specificity.

## Introduction

Mutualistic host-microbe interactions, or symbioses, can have profound impacts on development, resource utilization, and environmental responses. In extreme cases of obligate interactions, the timing and coordination of symbiosis establishment is essential for the evolution and persistence of the relationship itself. Many obligate microbial symbionts are vertically transmitted from parents to offspring, which stabilizes the relationship over evolutionary time. However, there are also obligate relationships that are re-established with each host generation through horizontal transmission where symbionts are acquired from the environment. These include many ecologically important interactions such as legume-rhizobia, plant-mycorrhizal fungi, and coral-algal symbioses. Therefore, how symbiosis is established and affects host development has been a key question in the study of host-microbe interactions (McFall-Ngai, 2002). Further, these relationships may be destabilized under environmental stress (Gupta et al., 2026; Helgoe et al., 2024; Li et al., 2026), so understanding the processes that maintain these interactions is essential to predicting their fate in a changing climate.

Corals form an obligate symbiosis with dinoflagellate algae in the family Symbiodiniaceae, where symbionts provide photosynthetic products to the host in exchange for inorganic nutrients, a high light environment, and shelter (Davy et al., 2012). The success of corals in nutrient-poor waters can be attributed to this mutualistic relationship, serving as the foundation for coral reef ecosystems (Muscatine and Porter, 1977). Despite the critical role of the symbionts in coral health, 85% of coral species acquire algae horizontally with each host generation during larval or juvenile life stages (Baird and Guest, 2009; Fadlallah, 1983). As symbiosis establishment occurs during a critical time of host metamorphosis and rapid growth, coordination between host and symbiont cell division may be important for successful development.

In cnidarian-algal symbioses, algae are initially acquired through the process of phagocytosis by cells in the host gastrodermal tissue, where symbiont cell division proceeds until populations reach a stable density (Colley and Trench, 1983; Schwarz et al., 1999). Previous studies on cell cycles during cnidarian-algal symbiosis establishment support host-symbiont cell cycle coordination and expansion of intracellular niche space (gastrodermal cells) (Gorman et al., 2025; Tivey et al., 2020a). However, these studies were performed in adult sea anemones, so the effect of symbiosis on host development was not explored. Only one study to date has investigated host-symbiont cell cycle dynamics in early life stage corals (Lecointe et al., 2016). Lecointe et al. (2016) found that symbiont cell division outpaced that of the host in larval and juvenile life stages, but that symbiont-to-host cell ratios were maintained, suggesting post-mitotic control of symbionts through expulsion or digestion. Given the importance of coral development, further studies are needed to evaluate the effect of symbiosis on early life stages at the cellular level. In addition, symbiosis establishment (Schnitzler et al., 2012) and algal cell division (Klueter et al., 2017) can be affected by environmental conditions, so understanding how symbiosis establishment is modulated by the environment, and how it affects host development is key to projecting the fate of corals under climate change.

Coral-algal symbioses are very sensitive to temperature. Climate change is causing an increase in the frequency and severity of marine heatwaves, which leads to the breakdown of the symbiotic relationship, known as coral bleaching. Repeated mass bleaching events have resulted in widespread coral mortality, reducing population density and leaving many coral species at risk of extinction (Hughes et al., 2018). Recovery of coral reefs therefore depends on survival and development of larval and juvenile offspring (Hughes and Tanner, 2000), but elevated temperature can also have a negative effect on symbiosis establishment. For example, exposure of larvae to thermal stress during symbiosis establishment reduces colonization rates and symbiont density, which leads to lower host survival and abnormal development (Kitchen et al., 2022; Schnitzler et al., 2012). Associating with a more thermally tolerant symbiont species could buffer these negative developmental effects, however the costs and benefits of host-symbiont interactions depend on host species, life stage, and environmental factors. For example, the algal species *Durusdinium trenchii* is considered thermally tolerant, which can protect corals from bleaching or dysbiosis (Berkelmans and van Oppen, 2006), but comes at an energetic cost to the host under ambient conditions (Matthews et al., 2017). Furthermore, in some corals such as *Acropora tenuis*, the dominant symbiont in adults can differ from that of early life stages (Abrego et al., 2009). Symbiont species also differ in their cell cycle dynamics (Tivey et al., 2020a), photosynthetic performance (Röser et al., 2024), and nutrient utilization (Wong et al., 2021), which could have differential effects on symbiosis establishment and host development.

In addition to functional differences, recognition mechanisms between host and symbiont could explain differences in colonization across algal species and environments. Although the precise interpartner dialogue that occurs between partners during symbiosis establishment has yet to be fully understood, there is evidence that microbe-associated molecular patterns (MAMP) on symbionts and host pattern recognition receptors (PRRs) are involved in host-symbiont recognition (reviewed in (Davy et al., 2012; Rosset et al., 2021)). The best studied MAMP-PRR system in cnidarian-algal symbiosis are lectin-glycan interactions (Davy et al., 2012). Glycans are complex carbohydrates that modify proteins and lipids and are involved in diverse processes such as cell communication, molecular function, and immunity. Host lectins recognize and bind glycan motifs on the cell surface of microbes, initiating immunotolerance or inflammatory responses (Varki, 2017). In cnidarian-algal symbiosis, removal or masking of symbiont glycans lowers symbiont uptake (Wood-Charlson et al., 2006) and proliferation (Bay et al., 2011), emphasizing their importance in symbiosis establishment. Previous studies have also found that the cell surface glycome varies across algal species (Tortorelli et al., 2022), and is modified by thermal stress (Maruyama et al., 2022). However, the responsiveness of algal glycomes has not yet been explored across different species and it is unknown whether glycan modifications mirror variation in physiological thermal tolerance, affecting symbiont uptake.

In this study, we experimentally manipulated algal symbionts by exposing them to heat stress prior to inoculation of larval hosts, to understand how elevated temperature affects interpartner recognition, symbiont acquisition and proliferation, and host development. We challenged larvae of the coral host, *Acropora aff. tenuis,* with four different algal species, including three native species (*Symbiodinium microadriaticum, Cladocopium goreaui, Durusdinium trenchii*) and one non-native species (*Breviolum minutum*), to leverage functional differences among partners and better understand host-symbiont specificity and cell cycle coordination. By measuring host-symbiont cell proliferation and characterizing the cell-surface glycome across species and temperatures, we sought to address the following questions: (1) Is host and symbiont cell division coordinated during colonization of coral larvae? (2) Do symbiont species differentially affect host development? and (3) How does thermal treatment affect symbiont recognition, uptake, and proliferation?

## Methods

### Coral spawning

Eight colonies of *Acropora* aff. *tenuis* (hereafter *A. tenuis* in this study) were collected from Sesoko Island, Okinawa, Japan (26°39.8440’ N; 127°52.4130’ E) and transported to Sesoko Marine Research Station, University of Ryukus. This species in Japan was previously referred to as *Acropora tenuis*, however it was recently shown to be a closely related, but distinct lineage (Bridge et al., 2023). Colonies were kept in running seawater with natural sunlight. On May 26th, 2023, all colonies spawned and egg/sperm bundles were collected. On spawning night, after fertilization, floating eggs were scooped with cups and transferred into new containers with filtered seawater (0.2 μm) twice to remove the remaining sperm. The seawater was changed daily until development into planula larvae.

### Experimental design

Four Symbiodiniaceae species were used for inoculation experiments: *Symbiodinium microadriaticum* (strain CCMP2458), *Breviolum minutum* (strain SSB01), *Cladocopium goreaui* (strain NIES2905), and *Durusdinium trenchii* (strain CCMP2556). Algal stocks were regularly maintained in f/2 media at 25°C with 12/12 light:dark cycle and transferred 1:10 every six weeks at Oregon State University in Oregon, USA. In 2023, cultures were transported to the Sesoko Marine Research Station, University of the Ryukyus for larval inoculations.

At the Sesoko Marine Research Station, each algal species was divided into two cultures and incubated under control (25°C) or elevated temperature (32°C) for three and a half days prior to inoculations. 1×10^7^ cells were subsampled from each replicate culture for glycan analysis. Six days post fertilization (dpf), larvae were inoculated with algae at a final concentration of 5×10^4^ cells/ml for 24 hours in 6-well plates, resulting in eight algal species by thermal pre-treatment combinations. All larvae were kept at 25°C at 75 umol/quanta/m^2^/sec PAR with water changes every other day. 10 larvae from two replicate wells (20 biological replicates) were sampled on day 3 and 9 post inoculation (dpi) to quantify host and symbiont cell proliferation.

To increase replication for glycan analysis and measure additional physiological information, algal thermal stress experiments in culture were repeated in 2025 at Oregon State University with three independent replicates per algal species and temperature treatment. Photosynthetic efficiency (Fv/Fm) was measured daily using a Walz Mini PAM-II fluorometer after one hour of dark acclimation using the following settings: Gain 3, Damping 2, Intensity 6. Algal density was quantified at the end of the 3.5 day experiment using an Invitrogen Countess Automated Cell Counter with three technical replicates per biological replicate to quantify any heat-stress induced mortality that occurred over the incubation period.

### Host cell proliferation, symbiont quantification, and colonization rate

Larvae were incubated in 10μM EdU (Click-iT EdU Alexa Fluor 555 imaging kit; Life Technologies, Eugene, OR, USA) for 24 h to label newly synthesized host DNA, a proxy for host cell proliferation. Samples were then fixed overnight at 4°C in 4% paraformaldehyde: 0.2% Tween in 1X PBS, permeabilized by washing with 0.2% Triton in 1X PBS (PBST) and blocked in 5% BSA. Incorporated EdU was then labelled with Alexa-Fluor 555 by incubating samples in Click-IT reaction buffer for 1 h. Reaction buffer was removed and host nuclei were stained with 0.2 μg/ml Hoechst 33342. Samples were stored in 87% glycerol at −20°C until confocal imaging.

Z stack images of six biological replicates of larvae per treatment were captured on the Zeiss LSM 780 NLO confocal microscope system in August 2023. EdU-labeled host nuclei were detected using 561nm excitation and 593nm emission wavelengths. Hoechst-labeled host nuclei were detected using 405nm excitation and 483nm emission and algal symbiont cells were identified using 633nm excitation and 684nm emission to capture chlorophyll a autofluorescence. In order to increase sample sizes within-algal species, twelve additional biological replicates were mounted and scored for the presence or absence of symbionts. Additional replicates inoculated with symbionts were imaged on a Leica confocal microscope system in July 2024 using the following detection settings: EdU: 553nm/570-620nm, Hoechst 405nm/410-490nm, Chla: 405nm/660-700nm excitation/emission. Larvae were imaged from the top to the midpoint of the Z axis. Samples with visible damage or abnormal development were excluded from downstream analysis (see <u>Table S</u>1 for final sample sizes).

Total area of EdU-labelling was measured from maximum projections generated in ImageJ by manually thresholding images to select EdU-positive signals after removing background noise using background subtraction. Individual Hoechst-stained nuclei could not be quantified in all samples due to background fluorescence. However, using a subset of well-stained, high resolution images, we found a positive correlation between the number of Hoechst-stained nuclei and total larval area (y=2.5x+2.7e5, R^2^=0.63, p=0.00026, <u>Fig S</u>1). Therefore we opted to normalize the EdU area by the total area of larvae measured using the polygon tool in the Hoechst channel. To determine symbiont density, symbionts in larvae were manually enumerated across stacks using the multi-point tool. Colonization rate was also quantified as the proportion of larvae with more than one symbiont out of the total number screened.

### Statistical analysis

Statistical analyses were performed in R v4.2.1. Due to its binary nature, colonization rates were compared across thermal treatments and symbiont species using a weighted generalized linear model. Continuous traits were evaluated for outliers, skewness, and normality. A square root and log10 transformation were applied to EdU and symbiont density *in hospite*, respectively to normalize distributions. Due to low colonization rates (<10%), larvae offered *C. goreaui* were excluded from subsequent analysis. A linear mixed effect model was used to evaluate the fixed effects of algal pre-treatment, species, timepoint, and their interaction on host cell division (EdU area), including confocal as a random effect to account for variance across systems (EdU ∼ algal pre-treatment*species*timepoint + (1|confocal) using the lme4 package (Bates et al., 2015). Confocal systems had a significant random effect on EdU but not symbiont density *in hospite*. Therefore symbiont density *in hospite* was evaluated using a linear model (symbiont density ∼ algal pre-treatment + species + timepoint). The effect of thermal pre-treatment and symbiont species on algal physiological traits in culture (Fv/Fm and cell density) was also tested using linear models from endpoint values (trait ∼ algal pre-treatment*species). The emmeans package (Estimated Marginal Means, aka Least-Squares Means) was used for post-hoc analyses of pairwise comparisons from linear models.

### Glycan analysis

Glycans were processed and quantified in two batches in 2023 and 2025 (<u>Table S</u>2). Due to low cell density in 2023, samples of *C. goreaui* were only included in the 2025 batch. For both batches, algal samples (∼ 1 × 10^7^ cells per sample) were pelleted at 3,100 x *g*, the supernatant decanted, and pellets frozen. Samples were thawed and washed sequentially twice in 1 mL 2X PBS, 1 ml Milli-Q water, and 50 mM ammonium bicarbonate, pelleting cells between washes (14,000 x *g* for 1 min). Cells were then treated with glycerol-free PNGase F (New England Biolabs, Rowley, MA, USA) under non-denaturing conditions following the manufacturer’s protocol to release N-glycans from cell surfaces. Pellets were incubated in 100 µL Glycobuffer 2, 900 µL ultrapure water and 3 µL PNGase F at 37°C for 72 h with occasional inversion. After incubation, a few drops of HCl were added to stop the reaction, and samples were frozen and subsequently freeze-dried for 24 h. The extracted *N-*glycans were cleaned using C18 solid-phase extraction (SPE) cartridges, with 5% acetic acid for elution. The eluted N-glycans were dried before reduction, which involved the addition of 10 μl of borane-ammonia complex at 10 μg/μl to each sample, followed by incubation in a water bath at 60°C for 1 h. Methanol was then added, and the samples were dried. This step was repeated four times to eliminate remaining borates. The reduced *N*-glycans were subsequently permethylated using a solid-phase method, as previously described (Fowowe et al., 2024; Kang et al., 2005). Briefly, sodium hydroxide beads stored in DMSO were loaded into an empty spin column and centrifuged at 1800 rpm for 2 min. The column was then rinsed with 200 μl DMSO and centrifuged again. *N*-glycan samples were reconstituted in 30 μl DMSO, 1.2 μl NaOH, and 20 μl iodomethane before being loaded to the prepared spin column. After incubation at room temperature for 25 min, another 20 μl of iodomethane was added, and the samples were incubated for an additional 15 min. The columns were then centrifuged at 1800 rpm for 2 min. Permethylated *N*-glycans were eluted with 30 μl acetonitrile (ACN) and dried. Prior to LC-MS/MS analysis, the samples were resuspended in 20% ACN containing 0.1% formic acid (FA) and centrifuged at 14800 rpm for 10 min.

LC-MS/MS analysis was performed using a Dionex 3000 Ultimate Nano-LC system (Thermo Fisher Scientific, Sunnyvale, CA, USA) coupled to Orbitrap-based mass spectrometers equipped with a nano-electrospray ionization (nano-ESI) source. The 2023 sample batch was analyzed using an Orbitrap Fusion Lumos Tribrid Mass Spectrometer (Thermo Fisher Scientific, San Jose, CA, USA), whereas the 2025 batch was analyzed using a Q Exactive HF Orbitrap mass spectrometer. Mobile phase A consisted of 2% ACN, 98% HPLC-grade water, and 0.1% FA, v/v, while mobile phase B consisted of 100% ACN with 0.1% FA, v/v. *N*-glycans were separated on a reversed-phase Acclaim PepMap C18 column (150 mm x 75 μm i.d.) maintained at 55°C, with a flow rate of 0.35 μl/min. A 60-min multistep gradient was used, beginning at 20% mobile phase B, increasing to 42% B, then to 55% B, and subsequently to 90% B before returning to 20% B for column equilibration.

Mass spectrometric data were acquired in positive ion mode using Orbitrap-based instruments. For both instruments, full MS scans were acquired at a resolution of 120,000 over a scan range of 400-2000 m/z, with an isolation width of 2 m/z. Data-dependent acquisition (DDA) was used for MS/MS fragmentation of the top 20 most intense precursor ions selected from each full MS scan. CID fragmentation was used on the Orbitrap Fusion Lumos Tribrid MS, whereas HCD fragmentation was used on the Q Exactive HF Orbitrap MS. Both acquisition methods used an AGC target of 1 × 10^5^. Raw data were processed with MultiGlycan software, as previously described (Hu, 2015). Peaks and retention times corresponding to charge states +1 to +4 were then manually checked in Xcalibur version 4.2 (Thermo Fisher Scientific). Glycan abundances were quantified in Skyline version 25.1.0.237 (MacCoss Lab, University of Washington).

To account for batch effects, we chose to analyze the data in the following ways: samples from the 2025 batch only (all species) and samples with species represented in both batches (*S. microadriaticum*, *B. minutum*, *D. trenchii*). For each data subset, half min gap filling was performed on absolute abundance count tables followed by probabilistic quotient normalization and log2 transformation. For the subset containing both batches, a batch correction was applied using limma (Ritchie et al., 2015) while preserving species and treatment level variation. Limma was then used to statistically evaluate differences in abundance of each glycan across species and treatment groups. As differential abundance of glycans was driven by species but not treatment in both datasets (File S2-S5, <u>Fig S</u>2), we opted to present differential abundance results for the 2025 batch only in order to be inclusive of all species. Glycan composition was evaluated at two levels of classification: (1) main structural types (high-mannose, hybrid, complex) and (2) subtypes representing specific glycan features (paucimannosides, sialylated (*N*-acetylneuramic acid NeuAc and N-glycolylneuramic acid NeuGc), fucosylated, sialofucosylated glycans). For ease of annotation, glycans were represented using a five-digit composition code corresponding to the numbers of *N*-acetylglucosamine, mannose and galactose combined, fucose, NeuAc, and NeuGc residues, respectively.

Relative abundance normalization was applied to all samples using individual glycan classification, structural categories, and subtypes. Compositional differences among species and treatments were visualized using NMDS plots and statistically evaluated by PERMANOVA using phyloseq (McMurdie and Holmes, 2013) and vegan (Dixon, 2003) packages.

## Results

### Thermal pre-treatment differentially affects algal physiology and survival but not larval colonization rate

After three days of experimental treatment, photosynthetic efficiency (Fv/Fm) of algal cultures was significantly reduced at elevated temperature compared to control (p<0.0001, <u>Table S</u>3). The magnitude of the response, however, differed by algal species (algal pre-treatment x species: p=0.0225, Fig 2A). Photosynthetic efficiency of *S. microadriaticum* and *B. minutum* was most sensitive to thermal stress, experiencing an 11.2-11.9% decline compared to control conditions. The sensitivity of these two species was significantly greater than *C. goreaui* and *D. trenchii*, where photosynthetic efficiency was reduced by 4.9% and 3.5% respectively. Further, photosynthetic efficiency of *D. trenchii* was not significantly affected by thermal stress (<u>Table S</u>4). Baseline photosynthetic efficiency in control conditions also differed between species, with *D. trenchii* exhibiting the highest baseline efficiency and *B. minutum* with the lowest (<u>Table S</u>5). Heat stress-induced mortality, measured as the change in final and initial algal cell density in culture between treatment groups, was only observed for *S. microadriaticum* (p=0.0045, <u>Fig S</u>3).

**Figure 1:**
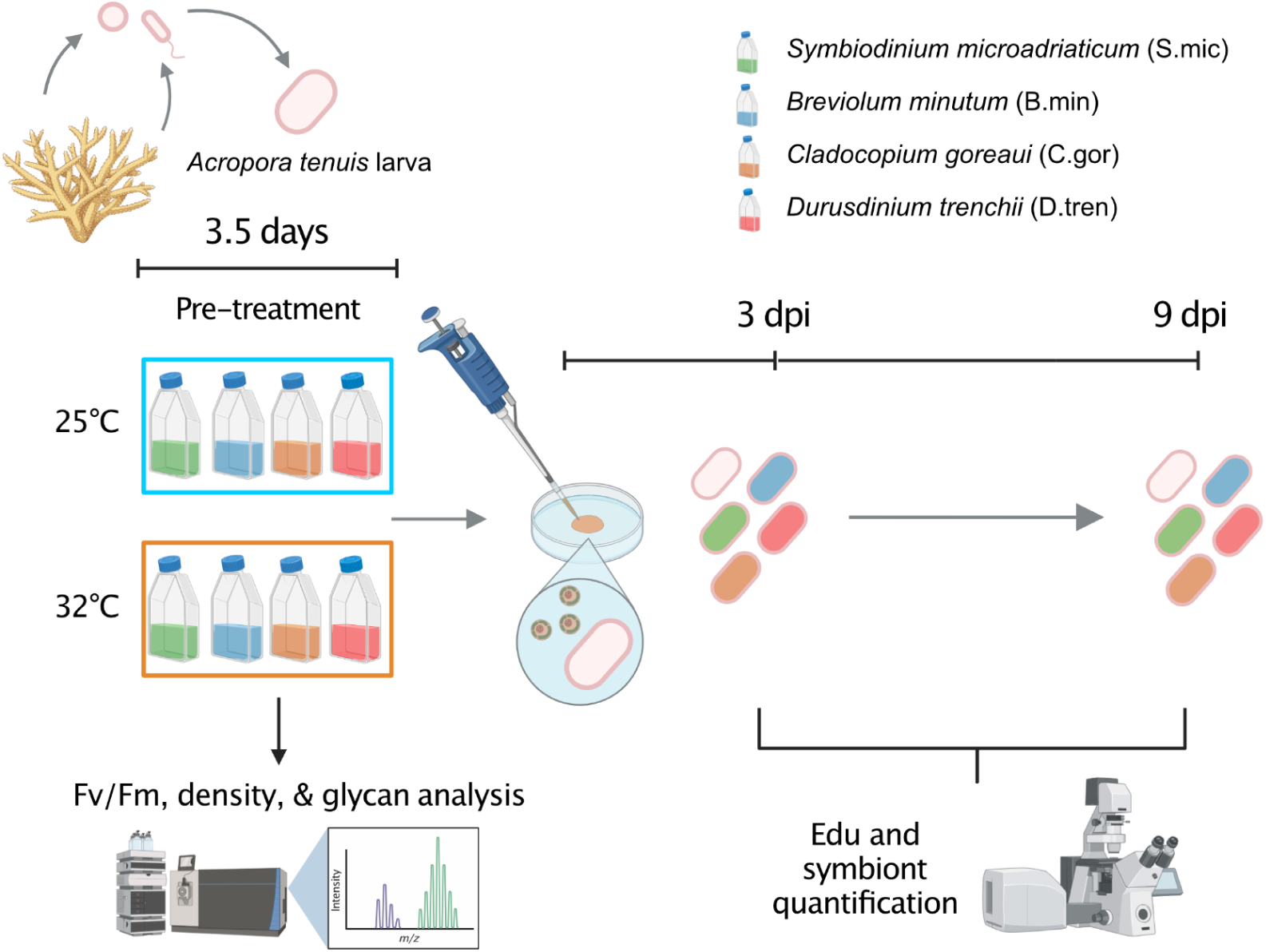
Experimental design. Four algal species, *Symbiodinium microadriaticum* (green), *Breviolum minutum* (blue), *Cladocopium goreaui* (orange), *Durusdinium trenchii* (red), were pre-treated in ambient or elevated temperature for 3.5 days before being offered to *Acropora tenuis* larvae. Symbiont samples for physiological and molecular analyses were taken during thermal pre-treatment and host samples were taken to measure for host proliferation (EdU staining) and symbiont quantification at days 3 and 9 post inoculation (dpi).

**Figure 2:**
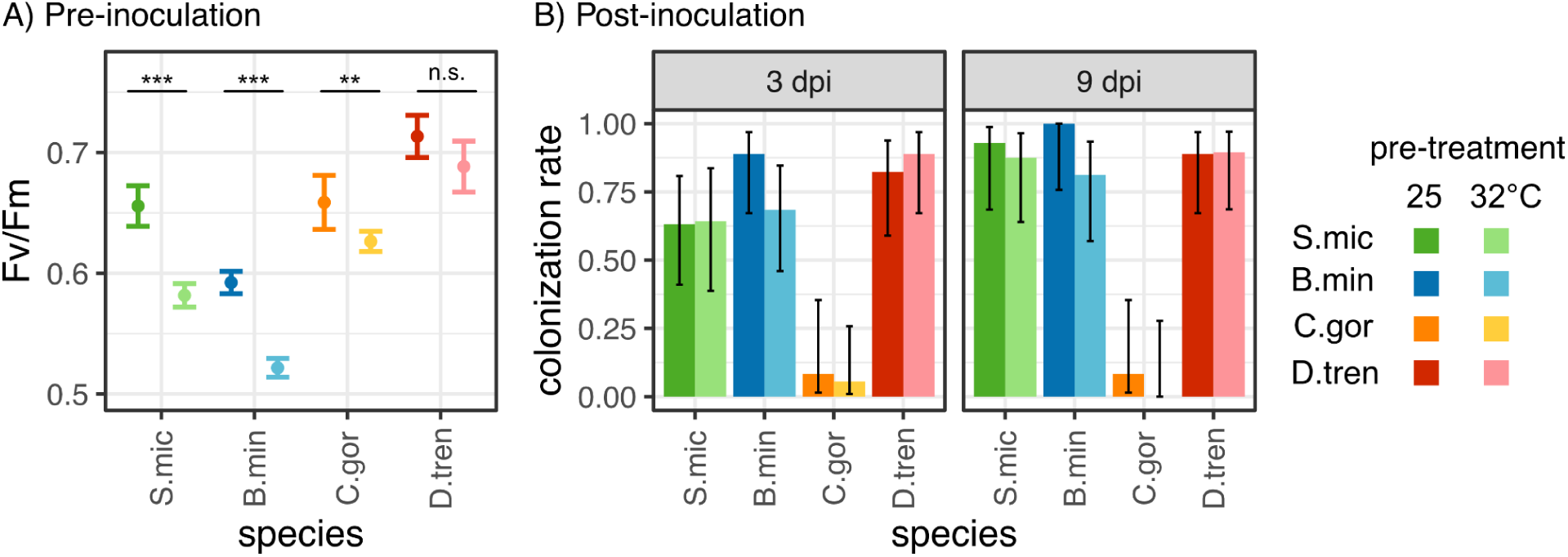
Algal photosynthetic efficiency and colonization rate of larvae following thermal pre-treatment of algae. A) Maximum quantum yield (Fv/Fm) of algae following 3 days of thermal treatment at 25°C and 32°C prior to larval inoculations. Error bars represent +/− 1 standard error around the mean from 3 biological replicates. Asterisks denote a significant effect of thermal treatment within each symbiont species (*: p<0.05, **:p<0.01, ***p<0.001). B) Colonization rate of coral larvae at 3 and 9 days post inoculation (dpi). Colonization rate was calculated as the proportion of larvae with symbionts divided by the total larvae sampled. Error bars represent 95% confidence intervals. For both panels, results are colored by species (S.mic: green, B.min: blue, C.gor: orange, D.tren: red) and shaded by algal pre-treatment (25°C: dark, 32°C: light).

*S. microadriaticum, B. minutum,* and *D. trenchii* all successfully colonized *A. tenuis* larvae, with greater than 77% of larvae colonized by day 9 post inoculation (Fig 2B). However, only 5.5% of *C. goreaui* treated larvae had symbionts, resulting in a significant effect of symbiont species on colonization rates (p<0.0001, <u>Table S</u>6). Colonization rates among symbiont species were not affected by thermal pre-treatment (p>0.05, <u>Table S</u>6).

### Thermal pre-treatment of algae negatively affects symbiont density across all species, but differentially affects host cell division

Thermal pre-treatment of algae reduced symbiont density in all species by 26% on average (p<0.001, <u>Table S</u>7), but did not affect symbiont proliferation over time (treatment x timepoint p>0.05, Fig 3A), suggesting that thermal pre-treatment affects initial density but not symbiont division. There was an overall effect of algal species on symbiont density, driven by slightly lower densities of *S. microadriaticum* compared to *B. minutum* and *D. trenchii* in hosts (<u>Table S</u>7). However, this pattern was not significant within pre-treatment groups or timepoints (<u>Table S</u>8), indicating symbiont densities did not vary greatly between the observed species.

**Figure 3:**
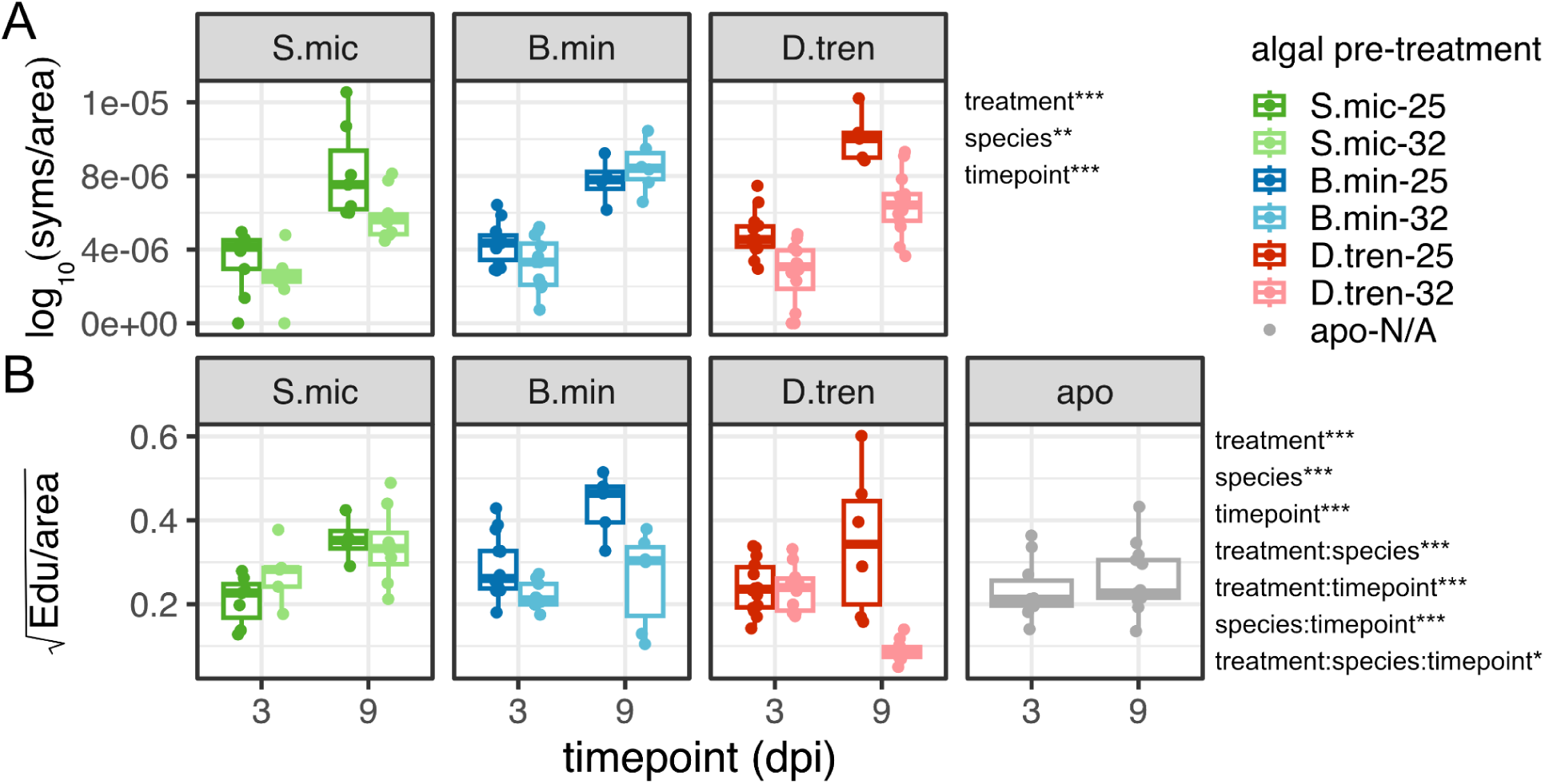
Symbiont density (A) and EdU-labelled host cell proliferation (B) over time across symbiont species and thermal pre-treatments. Panels and colors represent different symbiont species (S.mic: green, B.min: blue, C.gor: orange, D.tren: red), which are shaded by algal pre-treatment (25°C: dark, 32°C: light). Significance of algal pre-treatment, algal species, timepoint, and interactions evaluated using linear models are denoted by asterisks (*: p<0.05, **:p<0.01, ***p<0.001).

In contrast, there was a significant fixed effect of thermal pre-treatment of algae on larval cell division (p=3.22e^-6^, <u>Table S</u>9). Overall, larval cell division was reduced 25% when inoculated with algae pre-exposed to thermal stress compared to control algae. This reduction however was primarily driven by *B. minutum* and *D. trenchii*. Thermal pre-treatment of *B. minutum* and *D. trenchii* negatively affected larval cell division compared to control algae (Fig 3B, <u>Table S</u>10), but there was no significant difference in larval cell division between *S. microadriaticum* thermal treatments (<u>Table S</u>10), resulting in a thermal pre-treatment by species interaction (p=0.001, Fig 3B, <u>Table S</u>9). Further, host cell division in larvae not offered algae (aposymbiotic) was 65% greater than those inoculated with pre-stressed *D. trenchii* on day 9, suggesting a significant cost to this association (Tukey post-hoc, p=0.000015, <u>Table S</u>11).

### Symbiont density is positively correlated with host cell division

Symbiont densities in larvae were positively correlated to host cell proliferation when algae were not thermally treated (p<0.001, Fig 4, <u>Table S</u>12). Following thermal pre-treatment, symbiont density of all but one species maintained a positive correlation with host cell proliferation (Fig 4A), suggesting cell cycle coordination was not affected by heat stress for most associations. However, density of thermally-treated *D. trenchii* was negatively correlated with host cell proliferation, where higher symbiont densities of thermally-treated *D. trenchii* reduced host cell proliferation (Fig 4AB, <u>Table S</u>13), resulting in a significant effect of symbiont species and thermal pre-treatment on host cell cycle coordination (p<0.01, <u>Table S</u>12).

**Figure 4:**
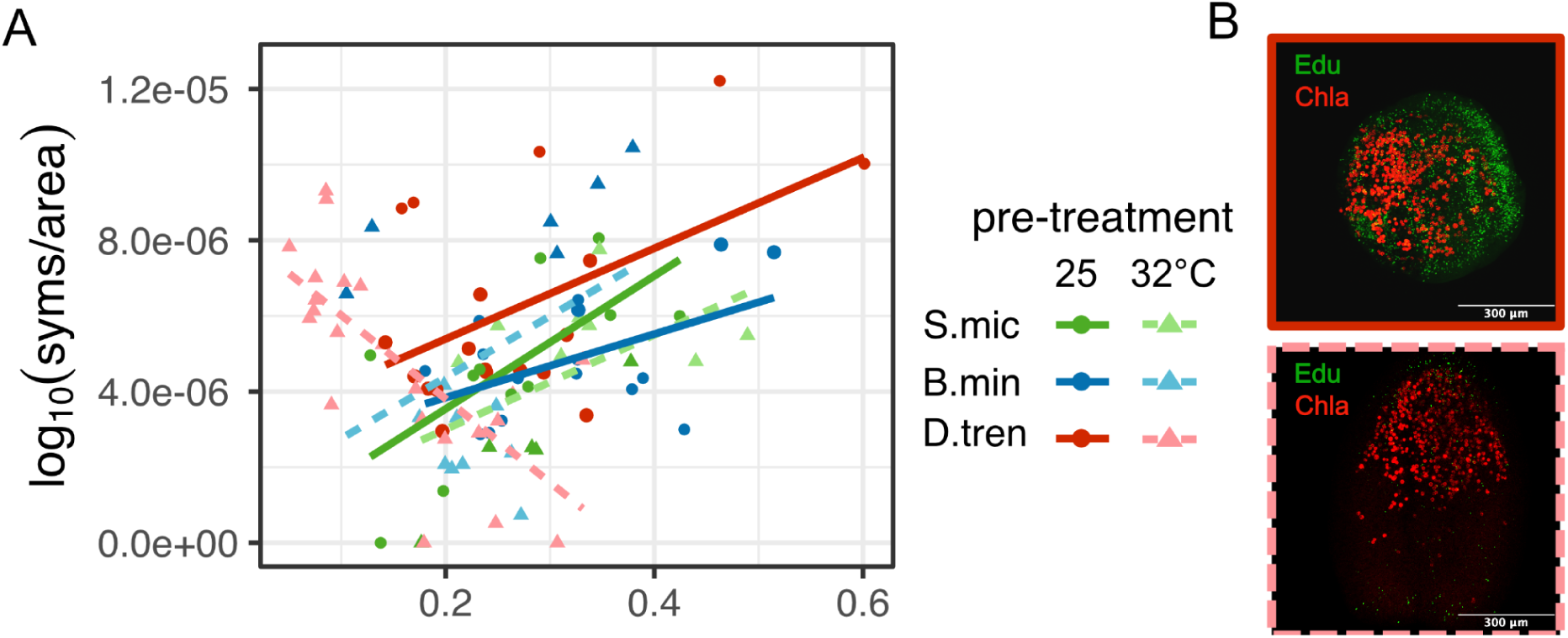
Correlation between larval cell proliferation (EdU) and symbiont density across symbiont species and thermal pre-treatments. A) Each point represents a single larva for which symbiont density and EdU was measured. Points are colored by symbiont species (*Symbiodinium microadriaticum* (green), *Breviolum minutum* (blue), *Durusdinium trenchii* (red)). Different shapes and shading indicate symbiont pre-treatment groups (circle: 25°C, triangle: 32°C). Lines indicate the correlation between traits within symbiont species-treatment groups (solid line: 25°C, dashed: 32°C). B) Maximum projection of confocal image showing dividing larval cells (EdU – green) and symbiont density (symbionts – red) for larvae with control (solid red) or thermally stressed (dashed pink) *D. trenchii*.

### Algal glycans vary across species but not thermal treatment

Thirty-three individual glycans were identified as consistent members of the algal glycome across experimental batches. Only one glycan type was exclusively detected in the elevated temperature condition (NeuGc fucosylated, glycan code 4-6-1-0-2), and two glycans were exclusively detected in the control condition (Sialylated NeuAc 8-6-0-1-0, NeuGc fucosylated 8-5-1-0-1). The presence of individual glycans was also largely consistent across symbiont species, with only one glycan (NeuGc fucosylated, 4-6-1-0-2) exclusively detected in *D. trenchii* symbionts. Of these 33 glycans, 7 were high-mannoside, 7 were hybrid, and 19 were complex glycan types. In all but one symbiont species, high-mannose was the predominant glycan conjugated to cell-surface proteins. However, in *B. minutum*, complex glycans made up the majority of the glycome (<u>Fig S</u>4).

Both glycan abundance and composition significantly differed across symbiont species, but not thermal treatments (Table <u>S</u>14-S15). Thirty percent of glycans (17/57) identified in batch 2 samples were differentially abundant across species (Fig 5B, <u>File S</u>2), including high-mannose, sialylated (NeuAc and NeuGc), fucosylated, sialofucosylated, and other complex glycans. *B. minutum* and *S. microadriaticum* had the greatest number of differentially abundant glycans (15), followed by *B. minutum* and *C. goreaui* (8), *S. microadriaticum* and *D. trenchii* (6), and *S. microadriaticum* and *C. goreaui* (4). Only one glycan significantly differed in abundance between *D. trenchii* and *B. minutum* or *C. goreaui*. Glycan composition also significantly differed between all species at the individual glycan level (Fig 5A, <u>Table S</u>16). At higher order levels (structural categories and glycan subtypes), however, compositional differences were primarily driven by *B. minutum* compared to all others, though glycan composition of *S. microadriaticum* and *D. trenchii* also significantly differed from one another at the structural and subtype level (Fig 5C, <u>Table S</u>16).

**Figure 5:**
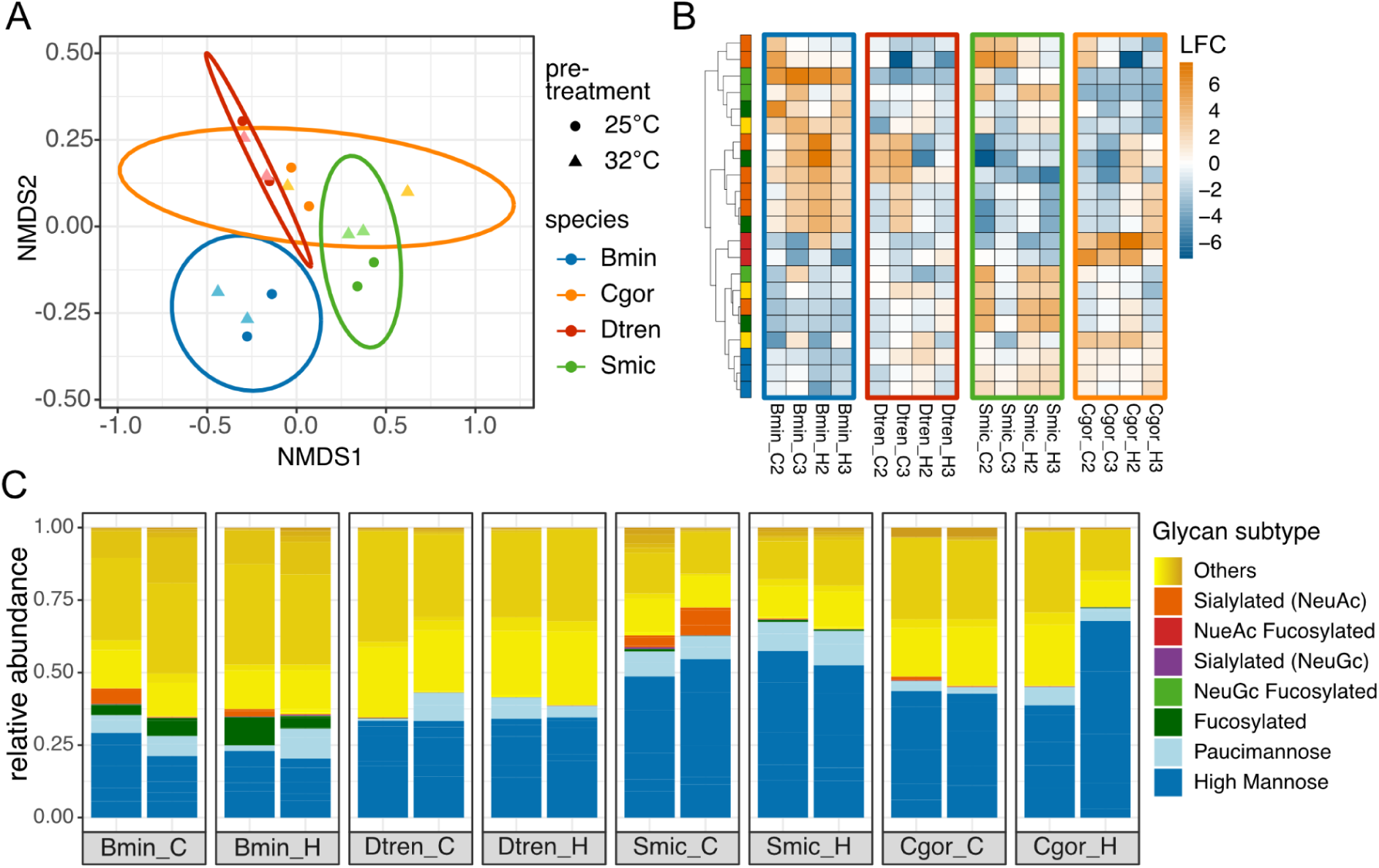
Symbiont glycan abundance and composition across species and treatment groups. (A) NMDS plot of glycan composition from 2025 samples. Colors represent symbiont species and shapes represent thermal treatment. (B) Heatmap of the 17 differentially abundant glycans (rows) across species for each sample (columns). Color shows log fold change relative to the mean abundance. Orange indicates higher abundance and blue indicates lower abundance. (C) Relative abundance of glycans for each sample. Each bar represents a single sample which is grouped by species and thermal treatment. Colors indicate relative abundance of glycan subtypes.

## Discussion

### Host and algal cell proliferation is coordinated during larval development

The establishment of cnidarian-algal endosymbiosis involves the uptake and rapid proliferation of symbionts within gastrodermal tissue. As the balance between host and symbiont cells is key to symbiosis stability (Falkowski et al., 1993; Muscatine, 1998; Rädecker et al., 2023), coordinated cell cycle regulation is required for successful establishment (Gorman et al., 2025; Xiang et al., 2020). Here we show that symbiont density is positively correlated to host cell proliferation under ambient conditions (Fig 3,4). This result is supported by previous studies in adult corals, which found that symbiont cell division is elevated during periods of rapid host growth such as the apical tips of branching corals (Gladfelter, 1983; Wilkerson et al., 1988). Further, in the vertically-transmitting coral *Montipora capitata*, symbiont density peaks during larval stages, which corresponds to a transition from nutrients provided by the parent to photosynthate from the symbiont (Huffmyer et al., 2025). On a cellular level, previous studies in the sea anemone Aiptasia have also found that symbiont colonization increases host cell proliferation (Gorman et al., 2022; Tivey et al., 2020a), consistent with results presented here. Although we did not track fitness metrics associated with increased host cell proliferation, other studies on coral symbiosis establishment suggest that acquisition of symbionts during larval stages improves post-settlement survival (Suzuki et al., 2013). Acquisition and proliferation of symbionts, therefore, are essential for coral development and reef recovery.

Cell cycle coordination between partners could be explained by cell signaling and/or nutrition. For example, a previous study on symbiosis establishment in Aiptasia observed localized, gastrodermal cell coordination around symbiont cells indicative of direct cell signaling between partners (Tivey et al., 2020a). However, epidermal cell division in Aiptasia is also upregulated during symbiosis establishment (Gorman et al., 2022; Tivey et al., 2020a). This could be explained by communication among host cells, where the presence of symbionts in the gastrodermis initiates growth signals to the epidermis. In addition, symbiont-derived nutrition during symbiosis establishment could increase the energy available for host cell division. Previous studies in coral larvae found that the quantity of photosynthetically-derived carbon was equivalent across host tissue layers despite localization of symbionts (Kopp et al., 2016). As we did not differentiate between tissue-level cell proliferation in this study, our results likely reflect increases in whole organism cell proliferation. Future studies of tissue-level microscopy, metabolomics, and gene expression studies would clarify the mechanisms of cell cycle coordination during symbiosis establishment.

### Thermally-treated algae differentially affect host cell proliferation

Some *Durusdinium* species are known to be thermally tolerant (Rowan, 2004), and can increase coral bleaching resistance by 1-2°C (Berkelmans and van Oppen, 2006). In our study, there was no significant effect of thermal pre-treatment on *D. trenchii* physiology (Fig 2A), supporting the high thermal tolerance previously observed in this species (Rowan, 2004). However, thermally-treated *D. trenchii* reduced host cell proliferation below that of control larvae and even below those not offered any symbionts (aposymbiotic), indicating a negative effect on host development. Crucially, this reduction in host cell proliferation did not coincide with a proportional reduction in symbiont density (Fig 3), suggesting thermally-treated *Durusdinium* are colonizing the host opportunistically.

These results are consistent with a growing body of literature supporting a tradeoff to hosting *D. trenchii*. *D. trenchii* symbionts translocate less carbon to their hosts compared to other symbiont species under ambient conditions (Cantin et al., 2009; Matthews et al., 2018), which results in reductions in coral growth and reproduction (Jones and Berkelmans, 2010; Jones and Berkelmans, 2011). Though the ability of *D. trenchii* to maintain carbon translocation under thermal stress may serve a fitness advantage at higher temperatures (Baker et al., 2013), we interestingly only observed a tradeoff to hosting *D. trenchii* following thermal pre-exposure. This discrepancy could be explained by inconsistent host and symbiont thermal history, as the host was not exposed to thermal stress during colonization. Corals with *D. trenchii* however also exhibit greater tissue loss and reduced growth under thermal stress despite apparently healthy symbiont densities and physiology (Chan et al., 2026), supporting the hypothesis that there is a cost to hosting *D. trenchii* even at elevated temperatures. Alternatively, the costs and benefits of hosting *D. trenchii* may be modulated by host species. For example, previous studies in *A. tenuis* found that corals with *Cladocopium* bleached less than those with *Durusdinium* (Abrego et al., 2008). Though these results are contrary to reports in other coral species (Berkelmans and van Oppen, 2006; Rowan, 2004; Silverstein et al., 2015) and thermal physiology of cultured algae shown here (Fig 2A), they are in support of a tradeoff to hosting thermally stressed *D. trenchii* in *A. tenuis* (Fig 3-4).

In contrast to the negative effects of thermally-treated *Durusdinium*, we found that larvae hosting thermally-treated *S. microadriaticum* maintained host cell proliferation (Fig 3), despite reductions in algal photosynthetic capacity and symbiont density (Fig 2-3), suggesting that this algal species continues to be beneficial for host development even under stress. This is supported by previous research in *A. tenuis*, where acquisition of a closely-related *Symbiodinium* species was associated with increased juvenile survival (Quigley et al., 2016; Suzuki et al., 2013). Elevated temperatures also did not affect early colonization of *A. tenuis* by *Symbiodinium* species in a prior study (Yorifuji et al., 2017), consistent with our results. Over time, however, long-term monitoring of *A. tenuis* juveniles at elevated temperatures in the same study showed that *Durusdinium* symbionts outcompete *Symbiodinium* (Yorifuji et al., 2017). Therefore, uptake of *S. microadriaticum* may provide an early developmental benefit to coral hosts that outweighs the negative effects of thermal stress, but community dynamics among symbiont species could constrain this benefit. Furthermore, *Durusdinium* symbionts tend to dominate *A. tenuis* juveniles when *Symbiodinium* is not present (Abrego et al., 2009). This coral species may therefore be particularly vulnerable to climate change if *D. trenchii* is able to opportunistically outcompete more beneficial symbionts at high temperatures during a critical time of host growth and development.

As symbiont communities of adult *A. tenuis* are dominated by *Cladocopium* symbionts (Abrego et al., 2009; Yamashita et al., 2014), we hypothesized that *Cladocopium* would increase host cell proliferation, thereby enhancing development and allowing *Cladocopium-*hosting corals to survive to adulthood. However, *C. goreaui* colonization rates of larvae were low in our study (Fig. 2). This is consistent with a prior study of *A. tenuis in situ*, which also found little-to-no acquisition of *Cladocopium* symbionts in month-old juveniles (Abrego et al., 2009). An alternative hypothesis for the transition to *Cladocopium-*dominated communities across life stages could be differences in microenvironments (Abrego et al., 2008). Larvae may be exposed to higher light and temperature environments in the water column compared to benthic life stages (Gleason and Wellington, 1995). Morphological complexity also increases in later life stages, which alters the light environment due to self shading (Voolstra et al., 2025). As *Cladocopium* symbionts are generally found in lower light and temperature environments (Rowan et al., 1997), competitive dynamics among symbionts or physiological benefits to the host could be different across life stages, driving the transition to *Cladocopium*-dominated adults. Although we cannot tease these processes apart in our study, we show that *C. goreaui* fails to colonize *A. tenuis* larvae, thereby indicating observed specificity in adults is not driven by enhanced larval development.

### Thermal pre-treatment negatively affects symbiont density in host larvae

Thermal pre-treatment of algae negatively affected initial symbiont density in larvae across all symbiont species, but did not affect symbiont proliferation after uptake (Figure 2B). This result indicates that thermal pre-treatment causes a reduction in the capacity of algae to colonize hosts or in the ability of the host to recognize or take up symbionts. Reduced symbiont density during thermal stress has been previously described in coral larvae, and leads to abnormal development and decreased survival (Kitchen et al., 2022; Schnitzler et al., 2012). However, both partners were exposed to thermal stress in these studies, so it is unclear whether decreases in symbiont density and host fitness are driven by the host, symbiont, or holobiont. When controlling for host and symbiont thermal exposure independently, a study in Aiptasia found that thermal pre-treatment of algae was responsible for decreases in initial symbiont density in hosts, as this effect was not observed when hosts alone were pre-treated with thermal stress and was equal to simultaneously exposing both partners (Kishimoto et al., 2020). This finding is supported by our study and indicates that modifications to algal physiology or biochemistry induced at high temperatures underlie reductions in symbiont density during uptake.

Reductions in photosynthetic performance of *S. microadriaticum*, *B. minutum*, and *C. goreaui* under elevated temperature are consistent with previous reports on the thermal sensitivity of these species (Berkelmans and van Oppen, 2006; Iglesias-Prieto et al., 1992; Russnak et al., 2021). As a decline in symbiont photophysiology can cause the breakdown of symbiosis, either due to reactive oxygen species production and/or nutritional dysregulation between partners (Helgoe et al., 2024; Lesser, 1996; Weis, 2008; Wooldridge, 2009), impaired photophysiology could lead to lower uptake during symbiosis establishment. However, symbiont density was reduced uniformly across symbiont species regardless of differences in photosynthetic efficiency, indicating that photosynthetic capacity does not explain differences in symbiont uptake in the present study. Furthermore, photosynthetic efficiency is only a proxy for photosynthesis. Additional studies using stable isotope tracer and metabolomics could assess the effects of thermal treatment on algal metabolism to better understand the role of nutrient exchange and specific metabolites in symbiont uptake. Alternatively, other physiological differences could be driving this response such as changes to algal motility. Motility is reduced under higher temperatures (Nitschke et al., 2015), which could impair the ability of the algae to come into contact with the host. Further studies quantifying how algal physiology and metabolism are affected by thermal stress across various symbiont species would clarify their importance in symbiosis establishment.

### Glycan composition does not explain differences in colonization between species or thermal treatments

There were no significant differences in glycan abundance or composition across thermal treatments, suggesting changes to the glycome do not underlie observed reductions in uptake density of thermally-treated algae. Glycan composition was instead driven by symbiont species (Fig 5). *S. microadriaticum*, *B. minutum* and *D. trenchii* however all had similar colonization rates and symbiont densities in larvae despite having differing glycan compositions. Further, the glycan profile of *C. goreaui*, which had a very low colonization rate, was almost identical to *D. trenchii*. Together, these results indicate that glycan composition on its own does not explain differences in symbiont colonization across species or thermal treatments.

Glycans have been previously identified as an important recognition molecule for symbiosis establishment. Enzymatic removal of glycans has been shown to reduce algal uptake (Wood-Charlson et al., 2006) and symbiont density (Bay et al., 2011), indicating that recognition of glycans by host lectins is an important step in symbiont colonization. In our study, we did not observe significant changes in glycan profiles with thermal stress, which could be a positive indication that symbiont recognition will not be affected in future oceans. This result is similar to a previous study on *B. minutum* glycomics, where only three, low abundance glycans differed across thermal treatments and the composition of major glycan types was unaffected (Maruyama et al., 2022). However, Maruyama et al. (2022) did identify a significant effect of thermal treatment when quantifying lectin binding of high-mannose and sialic acid via flow cytometry. The enzymatic extraction method used in our study (PNGase F) only cleaves glycans from N-linked glycoproteins, so it is possible that other glycan moieties, such as O-linked glycoproteins and glycolipids are changing in response to heat stress and driving differences in symbiont uptake across thermal treatments. Quantification of lectin binding, symbiont physiology, and nutritional exchange are necessary to tease apart these potential drivers of thermally-induced reductions in symbiont density.

Although differences in glycan abundance and composition were detected across species, they did not explain differences in colonization rate or symbiont density, suggesting that these glycans are not involved in discriminating between symbionts in *A. tenuis*. Glycan abundance has been previously reported to differ among species, but targeted manipulations of these differentially abundant glycans did not result in differences in uptake between compatible symbionts (Tortorelli et al., 2022). This supports the idea that differential abundance of glycans does not affect symbiont uptake or specificity, even of non-native symbionts such as *B. minutum* tested here. Rather, glycans common among symbiont species may act as broad molecular patterns that initiate phagocytosis. High-mannose glycans made up 30-70% of symbiont glycomes in this study, consistent with previous reports of 52-80% (Maruyama et al., 2022; Tivey et al., 2020b) and highlighting their importance in symbiont uptake. Two mannose-binding lectins have been isolated from corals, Millectin (Kvennefors et al., 2008) and PdC lectin (Vidal-Dupiol et al., 2009), and are putatively involved in symbiont uptake. These lectins however have extensive sequence diversity and multiple binding motifs, which confer broad binding of symbionts and bacterial pathogens alike (Kvennefors et al., 2008). As most coral lectins remain uncharacterized and there is a limited understanding of the specificity of glycan-lectin interactions, future research should explore the dynamics of glycan-lectin interactions among different host-symbiont combinations to determine the role of glycans in symbiont discrimination.

### Conclusion

Successful coral development depends in part on the timely establishment of symbiosis. Although corals with horizontal transmission can acquire symbionts during both larval and post-settlement juvenile stages, we show that symbiont proliferation in larval stages proportionally increases host cell division, suggesting that early acquisition of symbionts is beneficial for host development. Although thermal pre-treatment of algae negatively affected symbiont densities in larvae, effects on host cell proliferation were species-specific and did not mirror differences in thermal tolerance of algae. These results indicate that the success of *A. tenuis* development depends on symbiotic partner, but a mismatch between environmental tolerance of algal species and developmental benefits may constrain reef recovery under climate change. Reductions in symbiont density under thermal stress do not seem to be driven by changes in glycan composition, suggesting physiology of symbionts, nutritional exchange, and/or other recognition mechanisms may therefore be more important in determining symbiont colonization and developmental outcomes in future oceans.

## Acknowledgements

We would like to thank Frederic Sinniger and Mori Jinza from Sesoko Marine Research Station, Univ. Rykyus for assisting with coral spawning. We would also like to thank Mark Phillips for his input on glycan analyses. Lastly, we would like to thank the OSU CQLS center, particularly Anne-Marie Girard-Pohjanpelto, for training and assistance on the confocal system. Coral colonies were collected under permits issued by the Okinawa Prefectural Government, Japan, to SH (No. 4-28).

## Data availability

Data and scripts to recreate analyses are publicly available on github (https://github.com/mruggeri55/Aten-larvae-dev). The raw spectrum files for the LC-MS/MS analysis are available on MassIVE database with ID: MSV000102430

