## Supplemental document 1 for "Thermal pre-treatment of algal symbiont species differentially affects coral development"

### Supplemental figures:

Figure S1. Hoechst to size correlation showing a significant positive relationship between the area of Hoechst-stained nuclei and total larvae area.

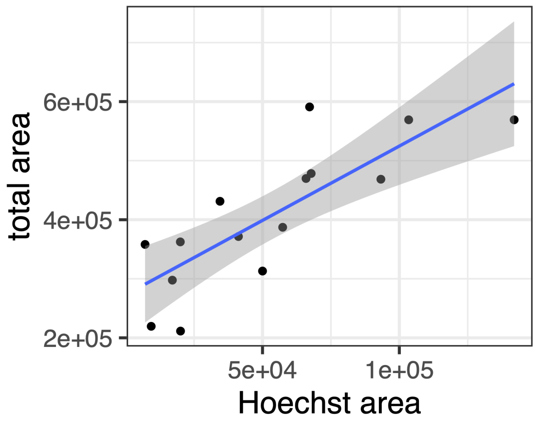


Figure S2. PCA of glycan composition from batch-corrected dataset showing separation across species but not treatment. Note, *Cladocopium goreaui* is not included in the batch corrected analysis as it was only included in the second batch. Therefore we opted to present results from batch 2 only to be inclusive of all species (see Fig 5 in main text).

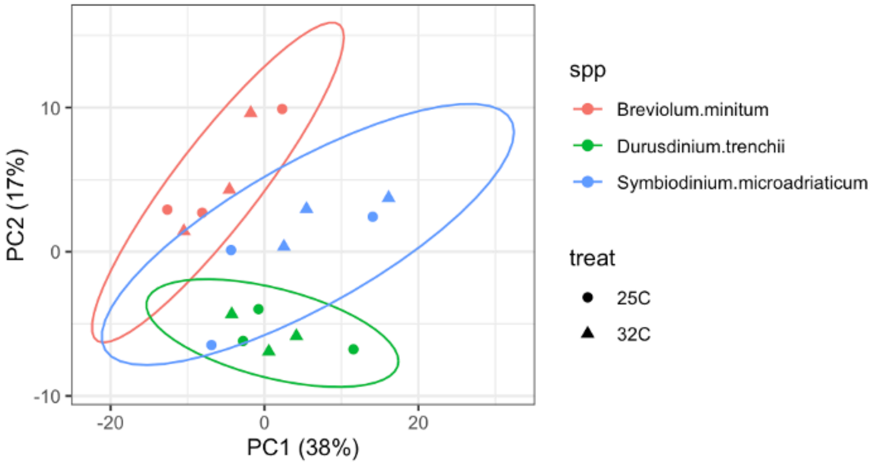


Figure S3. Change in algal density in culture before and after 3 days of thermal treatment (blue=25C, orange=32C).

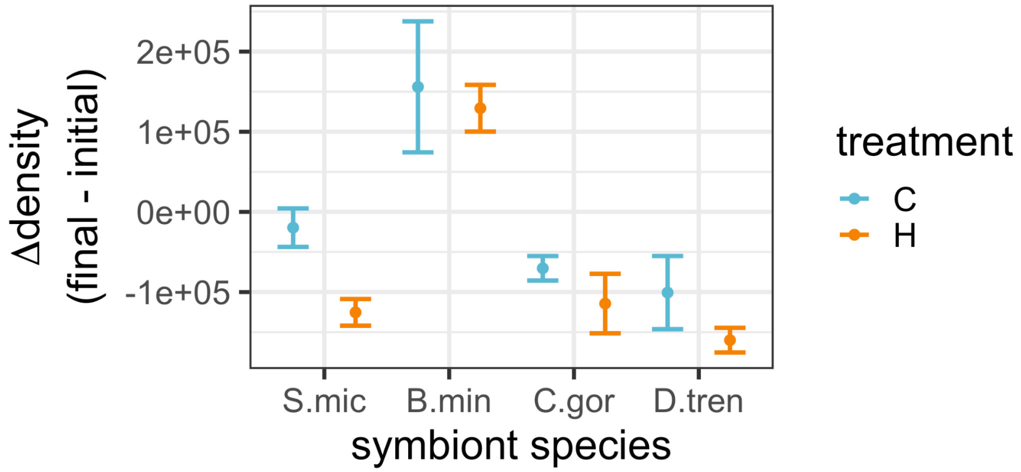


Figure S4. Relative abundance of glycans based on higher order classifications. C: complex, H: hybrid, M: mannose.

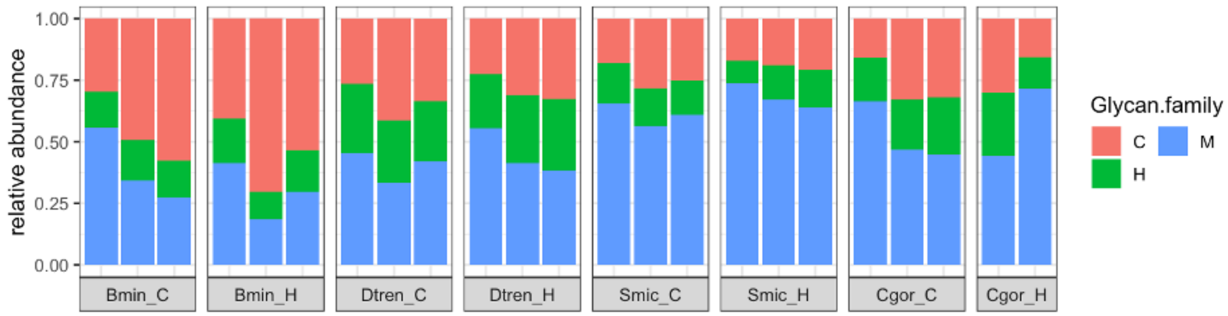

### Supplemental Tables:

Table S1. Sample sizes for confocal images after quality control.

| species | treatment | timepoint | n |
|---|---|---|---|
| apo | NA | 3 | 13 |
| apo | NA | 9 | 16 |
| Smic | 25 | 3 | 9 |
| Smic | 25 | 9 | 7 |
| Smic | 32 | 3 | 9 |
| Smic | 32 | 9 | 10 |
| Bmin | 25 | 3 | 16 |
| Bmin | 25 | 9 | 7 |
| Bmin | 32 | 3 | 13 |
| Bmin | 32 | 9 | 7 |
| Cgor | 25 | 3 | 1 |
| Cgor | 25 | 9 | 1 |
| Cgor | 32 | 3 | 1 |
| Dtren | 25 | 3 | 14 |
| Dtren | 25 | 9 | 6 |
| Dtren | 32 | 3 | 12 |
| Dtren | 32 | 9 | 13 |

Table S2. Algal glycan sample metadata.

| sample | spp | treat | experimental batch | quantification batch |
|---|---|---|---|---|
| Breviolum.minitum_25C_1 | Bmin | C | 2023 | A |
| Breviolum.minitum_25C_2 | Bmin | C | 2025 | B |
| Breviolum.minitum_25C_3 | Bmin | C | 2025 | B |
| Breviolum.minitum_32C_1 | Bmin | H | 2023 | A |
| Breviolum.minitum_32C_2 | Bmin | H | 2025 | B |
| Breviolum.minitum_32C_3 | Bmin | H | 2025 | B |
| Durusdinium.trenchii_25C_1 | Dtren | C | 2023 | A |
| Durusdinium.trenchii_25C_2 | Dtren | C | 2025 | B |
| Durusdinium.trenchii_25C_3 | Dtren | C | 2025 | B |
| Durusdinium.trenchii_32C_1 | Dtren | H | 2023 | A |
| Durusdinium.trenchii_32C_2 | Dtren | H | 2025 | B |
| Durusdinium.trenchii_32C_3 | Dtren | H | 2025 | B |
| Symbiodinium.microadriaticum_25C_1 | Smic | C | 2023 | A |
| Symbiodinium.microadriaticum_25C_2 | Smic | C | 2025 | B |
| Symbiodinium.microadriaticum_25C_3 | Smic | C | 2025 | B |
| Symbiodinium.microadriaticum_32C_1 | Smic | H | 2023 | A |
| Symbiodinium.microadriaticum_32C_2 | Smic | H | 2025 | B |
| Symbiodinium.microadriaticum_32C_3 | Smic | H | 2025 | B |
| Cladocopium.goreaui_25C_1 | Cgor | C | N/A | C |
| Cladocopium.goreaui_25C_2 | Cgor | C | 2025 | B |
| Cladocopium.goreaui_25C_3 | Cgor | C | 2025 | B |
| Cladocopium.goreaui_32C_1 | Cgor | H | 2025 | B |
| Cladocopium.goreaui_32C_2 | Cgor | H | 2025 | B |

Table S3. Linear model results testing the effect of thermal treatment, species, and their interaction on photosynthetic efficiency (Fv/Fm).

| model term | df1 | df2 | F.ratio | p.value |
|---|---|---|---|---|
| treatment | 1 | 16 | 66.468 | <.0001 |
| spp | 3 | 16 | 92.358 | <.0001 |
| treatment:spp | 3 | 16 | 4.212 | 0.0225 |

Table S4. Within-species treatment effect on photosynthetic efficiency (Fv/Fm).

| Species | contrast | estimate | SE | df | t.ratio | p.value |
|---|---|---|---|---|---|---|
| Smic | C - H | 0.074 | 0.0124 | 16 | 5.973 | <.0001 |
| Bmin | C - H | 0.0707 | 0.0124 | 16 | 5.704 | <.0001 |
| Cgor | C - H | 0.0323 | 0.0124 | 16 | 2.61 | 0.019 |
| Dtren | C - H | 0.025 | 0.0124 | 16 | 2.018 | 0.0607 |

Table S5. Within-treatment species effect on photosynthetic efficiency (Fv/Fm).

| Treatment | contrast | estimate | SE | df | t.ratio | p.value |
|---|---|---|---|---|---|---|
| Control | A - B | 0.0633 | 0.0124 | 16 | 5.112 | 0.0005 |
| Control | A - C | -0.003 | 0.0124 | 16 | -0.242 | 0.9948 |
| Control | A - D | -0.0577 | 0.0124 | 16 | -4.655 | 0.0014 |
| Control | B - C | -0.0663 | 0.0124 | 16 | -5.354 | 0.0003 |
| Control | B - D | -0.121 | 0.0124 | 16 | -9.767 | <.0001 |
| Control | C - D | -0.0547 | 0.0124 | 16 | -4.413 | 0.0022 |
| Heat | A - B | 0.06 | 0.0124 | 16 | 4.843 | 0.0009 |
| Heat | A - C | -0.0447 | 0.0124 | 16 | -3.606 | 0.0114 |
| Heat | A - D | -0.1067 | 0.0124 | 16 | -8.61 | <.0001 |
| Heat | B - C | -0.1047 | 0.0124 | 16 | -8.449 | <.0001 |
| Heat | B - D | -0.1667 | 0.0124 | 16 | -13.453 | <.0001 |
| Heat | C - D | -0.062 | 0.0124 | 16 | -5.005 | 0.0007 |

Table S6. Weighted binomial GLM results testing the effects of algal pre-treatment, species, and timepoint on colonization rates.

| *Predictors* | *Odds Ratios* | *CI* | *p* |
|---|---|---|---|
| (Intercept) | 1.78 | 0.71 – 4.68 | 0.227 |
| symTreat [32] | 0.65 | 0.31 – 1.34 | 0.249 |
| symClade [B] | 1.66 | 0.69 – 4.13 | 0.261 |
| symClade [C] | 0.02 | 0.00 – 0.06 | **<0.001** |
| symClade [D] | 2.22 | 0.90 – 5.78 | 0.090 |
| timepoint | 1.15 | 1.02 – 1.31 | **0.024** |
| Observations | 16 | | |

Table S7. Linear model results of log10 transformed symbiont density in larvae based on symbiont pre-treatment (symTreat), species (symClade), and timepoint.

| | sym.count.log/size | | |
|---|---|---|---|
| *Predictors* | *Estimates* | *CI* | *p* |
| (Intercept) | 0.0000037 | 0.0000030 – 0.0000043 | **<0.001** |
| symTreat [32] | -0.0000017 | -0.0000023 – -0.0000011 | **<0.001** |
| symClade [B] | 0.0000010 | 0.0000003 – 0.0000018 | **0.008** |
| symClade [D] | 0.0000009 | 0.0000002 – 0.0000016 | **0.015** |
| timepoint [9] | 0.0000042 | 0.0000036 – 0.0000048 | **<0.001** |
| Observations | 115 | | |
| $R^2$ / $R^2$ adjusted | 0.647 / 0.634 | | |

Table S8. Pairwise differences of symbiont density across symbiont species and treatments.

| contrast | estimate | SE | df | t.ratio | p.value |
|---|---|---|---|---|---|
| A symTreat25 - B symTreat25 | -1.02E-06 | 3.75E-07 | 110 | -2.7228 | 0.0788 |
| A symTreat25 - D symTreat25 | -8.74E-07 | 3.54E-07 | 110 | -2.4732 | 0.1412 |
| **A symTreat25 - A symTreat32** | **1.66E-06** | **2.96E-07** | **110** | **5.6284** | **0.0000** |
| A symTreat25 - B symTreat32 | 6.44E-07 | 4.79E-07 | 110 | 1.3450 | 0.7592 |
| A symTreat25 - D symTreat32 | 7.90E-07 | 4.54E-07 | 110 | 1.7423 | 0.5072 |
| B symTreat25 - D symTreat25 | 1.46E-07 | 3.52E-07 | 110 | 0.4151 | 0.9984 |
| B symTreat25 - A symTreat32 | 2.69E-06 | 4.76E-07 | 110 | 5.6399 | **0.0000** |
| **B symTreat25 - B symTreat32** | **1.66E-06** | **2.96E-07** | **110** | **5.6284** | **0.0000** |
| B symTreat25 - D symTreat32 | 1.81E-06 | 4.51E-07 | 110 | 4.0132 | **0.0015** |
| D symTreat25 - A symTreat32 | 2.54E-06 | 4.68E-07 | 110 | 5.4233 | **0.0000** |
| D symTreat25 - B symTreat32 | 1.52E-06 | 4.69E-07 | 110 | 3.2399 | **0.0192** |
| **D symTreat25 - D symTreat32** | **1.66E-06** | **2.96E-07** | **110** | **5.6284** | **0.0000** |
| A symTreat32 - B symTreat32 | -1.02E-06 | 3.75E-07 | 110 | -2.7228 | 0.0788 |
| A symTreat32 - D symTreat32 | -8.74E-07 | 3.54E-07 | 110 | -2.4732 | 0.1412 |
| B symTreat32 - D symTreat32 | 1.46E-07 | 3.52E-07 | 110 | 0.4151 | 0.9984 |

Table S9. Linear mixed effect model results testing the effect of symbiont pre-treatment (symTreat), species (symClade), timepoint, and their interaction on square root transformed Edu values, with confocal system as a random effect.

| *Row* | *Sum Sq* | *Mean Sq* | *NumDF* | *DenDF* | *F value* | *Pr(>F)* |
|---|---|---|---|---|---|---|
| symTreat | 0.1331 | 0.1331 | 1 | 87.1112 | 24.7722 | 0.0000 |
| symClade | 0.0954 | 0.0477 | 2 | 87.1524 | 8.8744 | 0.0003 |
| timepoint | 0.0778 | 0.0778 | 1 | 87.0201 | 14.4891 | 0.0003 |
| symTreat:symClade | 0.0801 | 0.0400 | 2 | 87.4267 | 7.4539 | 0.0010 |
| symTreat:timepoint | 0.1091 | 0.1091 | 1 | 87.0281 | 20.3080 | 0.0000 |
| symClade:timepoint | 0.0813 | 0.0406 | 2 | 87.2060 | 7.5631 | 0.0009 |
| symTreat:symClade:timepoint | 0.0426 | 0.0213 | 2 | 87.1674 | 3.9639 | 0.0225 |

| *Row* | *npar* | *logLik* | *AIC* | *LRT* | *Df* | *Pr(>Chisq)* |
|---|---|---|---|---|---|---|
| | 14 | 91.8267 | -155.6534 | NA | NA | NA |
| (1 \| confocal) | 13 | 89.5420 | -153.0839 | 4.5695 | 1 | 0.0325 |

Table S10. Pairwise comparisons between symbiont species and pre-treatment (symTreat) groups on host Edu derived from linear mixed effect model described in Table S9.

| **contrast** | **estimate** | **SE** | **df** | **t.ratio** | **p.value** |
|---|---|---|---|---|---|
| A symTreat25 - B symTreat25 | -0.0717477 | 0.03022688 | 87.3422244 | -2.3736387 | 0.177031 |
| A symTreat25 - D symTreat25 | -0.0130222 | 0.02912138 | 87.0114754 | -0.44717 | 0.99767497 |
| **A symTreat25 - A symTreat32** | -0.0116107 | 0.03162685 | 87.4990389 | -0.3671164 | 0.99909971 |
| A symTreat25 - B symTreat32 | 0.04134111 | 0.03001291 | 87.0006139 | 1.37744458 | 0.74029592 |
| A symTreat25 - D symTreat32 | 0.12337542 | 0.0278708 | 87.0584527 | 4.42669047 | **0.0003879** |
| B symTreat25 - D symTreat25 | 0.05872548 | 0.02671458 | 87.5557987 | 2.19825569 | 0.24914091 |
| B symTreat25 - A symTreat32 | 0.06013696 | 0.02833665 | 87.0372069 | 2.12223262 | 0.28573869 |
| **B symTreat25 - B symTreat32** | **0.11308881** | **0.02752103** | **87.3762764** | **4.10917805** | **0.00122202** |
| B symTreat25 - D symTreat32 | 0.19512311 | 0.02488753 | 87.209865 | 7.84019649 | **4.73E-10** |
| D symTreat25 - A symTreat32 | 0.00141148 | 0.02832697 | 87.7193613 | 0.04982794 | 0.99999996 |
| D symTreat25 - B symTreat32 | 0.05436332 | 0.0263404 | 87.0214535 | 2.06387588 | 0.31586827 |
| **D symTreat25 - D symTreat32** | **0.13639763** | **0.02392061** | **87.1649967** | **5.70209732** | **2.34E-06** |
| A symTreat32 - B symTreat32 | 0.05295185 | 0.02904285 | 87.5457219 | 1.8232319 | 0.4564562 |
| **A symTreat32 - D symTreat32** | **0.13498616** | **0.02648211** | **87.3809755** | **5.09725818** | **2.86E-05** |
| **B symTreat32 - D symTreat32** | **0.08203431** | **0.02493226** | **87.0579105** | **3.29028789** | **0.0175597** |

Table S11. Pairwise comparisons (Tukey post hoc) of Edu between larvae offered no symbionts (apo symTreatNA) or symbiont species pre-treated at 32C (symTreat32) on 9 days post inoculation.

| comparison | diff | lwr | upr | p adj |
|---|---|---|---|---|
| A symTreat32 - apo symTreatNA | 0.08067303 | -0.0115813 | 0.17292737 | 0.10433708 |
| B symTreat32 - apo symTreatNA | 0.00246526 | -0.0988613 | 0.10379183 | 0.99989656 |
| **D symTreat32 - apo symTreatNA** | **-0.1701752** | **-0.2523619** | **-0.0879885** | **1.58E-05** |
| B symTreat32 - A symTreat32 | -0.0782078 | -0.1890837 | 0.03266816 | 0.24559893 |
| **D symTreat32 - A symTreat32** | **-0.2508482** | **-0.3445554** | **-0.1571409** | **1.17E-07** |
| **D symTreat32 - B symTreat32** | **-0.1726404** | **-0.2752916** | **-0.0699893** | **0.00036219** |

Table S12. ANOVA results from a linear model testing the effects of symbiont species and thermal pre-treatment on the relationship between symbiont density and host cell proliferation.

| Term | Df | Sum Sq | Mean Sq | F value | Pr(>F) |
|---|---|---|---|---|---|
| sqrt_Edu_size | 1 | 1.88E-11 | 1.88E-11 | 4.12034233 | 0.0455354 |
| symClade | 2 | 3.04E-11 | 1.52E-11 | 3.32246374 | 0.04086041 |
| symTreat | 1 | 1.44E-12 | 1.44E-12 | 0.3144877 | 0.57643216 |
| sqrt_Edu_size:symClade | 2 | 2.53E-11 | 1.27E-11 | 2.77334251 | 0.06817897 |
| sqrt_Edu_size:symTreat | 1 | 2.84E-11 | 2.84E-11 | 6.22451577 | 0.01455734 |
| symClade:symTreat | 2 | 4.40E-11 | 2.20E-11 | 4.81212162 | 0.01050559 |
| sqrt_Edu_size:symClade:symTreat | 2 | 6.38E-11 | 3.19E-11 | 6.9876813 | 0.00155891 |
| Residuals | 84 | 3.84E-10 | 4.57E-12 | NA | NA |

Table S13. Correlation coefficients and significance of symbiont density and host cell proliferation for each symbiont species and thermal treatment combination.

| symClade | symTreat | sqrt_Edu_size.trend | SE | df | t.ratio | p.value |
|---|---|---|---|---|---|---|
| A | 25 | 1.76E-05 | 7.40E-06 | 84 | 2.37828845 | 0.01966208 |
| B | 25 | 8.41E-06 | 5.85E-06 | 84 | 1.43850707 | 0.1540058 |
| D | 25 | 1.20E-05 | 4.43E-06 | 84 | 2.70273752 | 0.00832039 |
| A | 32 | 1.25E-05 | 7.02E-06 | 84 | 1.77844045 | 0.07894977 |
| B | 32 | 1.62E-05 | 7.51E-06 | 84 | 2.15486569 | 0.03403502 |
| D | 32 | -2.21E-05 | 5.48E-06 | 84 | -4.0367939 | 0.00011912 |

Table S14. PERMANOVA results testing the effect of algal species on glycan composition (relative abundance) across categorical levels: A) individual glycans, B) glycan types, C) glycan families.

A) Individual glycans

| | Df | SumOfSqs | R2 | F | Pr(>F) |
|---|---|---|---|---|---|
| **species** | 3 | 0.72607 | 0.44941 | 5.1695 | 0.001 |
| **Residual** | 19 | 0.88954 | 0.55059 | | |
| **Total** | 22 | 1.61562 | 1 | | |

B) Glycan type

| | Df | SumOfSqs | R2 | F | Pr(>F) |
|---|---|---|---|---|---|
| **species** | 3 | 0.407 | 0.46623 | 5.532 | 0.001 |
| **Residual** | 19 | 0.46596 | 0.53377 | | |
| **Total** | 22 | 0.87296 | 1 | | |

C) Glycan family

| | Df | SumOfSqs | R2 | F | Pr(>F) |
|---|---|---|---|---|---|
| **species** | 3 | 0.33757 | 0.60713 | 9.7872 | 0.002 |
| **Residual** | 19 | 0.21844 | 0.39287 | | |
| **Total** | 22 | 0.55601 | 1 | | |

Table S15. PERMANOVA results testing the effect of algal pre-treatment on glycan composition (relative abundance) across categorical levels: A) individual glycans, B) glycan types, C) glycan families.

A) Individual glycans

| | Df | SumOfSqs | R2 | F | Pr(>F) |
|---|---|---|---|---|---|
| **treatment** | 1 | 0.00007 | 0.00004 | 9.00E-04 | 0.996 |
| **Residual** | 21 | 1.61555 | 0.99996 | | |
| **Total** | 22 | 1.61562 | 1 | | |

B) Minor glycan type

| | Df | SumOfSqs | R2 | F | Pr(>F) |
|---|---|---|---|---|---|
| **treatment** | 1 | 0.0022 | 0.00251 | 0.0529 | 0.977 |
| **Residual** | 21 | 0.87076 | 0.99749 | | |
| **Total** | 22 | 0.87296 | 1 | | |

C) Major glycan type (mannose, hybrid, complex)

| | Df | SumOfSqs | R2 | F | Pr(>F) |
|---|---|---|---|---|---|
| **treat** | 1 | -0.00116 | -0.00208 | -0.0436 | 0.997 |
| **Residual** | 21 | 0.55717 | 1.00208 | | |
| **Total** | 22 | 0.55601 | 1 | | |

Table S16. Pairwise PERMANOVA results testing whether glycan composition (relative abundance) differs between algal species across categorical levels: A) individual glycans, B) glycan types, C) glycan families.

A) Individual glycans

| species 1 | species 2 | F.Model | R2 | p.value | p.adjusted |
|---|---|---|---|---|---|
| Bmin | Dtren | 5.061984 | 0.3360768 | 0.003 | 0.003 |
| Bmin | Smic | 4.811256 | 0.3248378 | 0.003 | 0.003 |
| Bmin | Cgor | 5.001941 | 0.357232 | 0.002 | 0.002 |
| Dtren | Smic | 7.435621 | 0.4264615 | 0.005 | 0.005 |
| Dtren | Cgor | 6.02595 | 0.4010362 | 0.009 | 0.009 |
| Smic | Cgor | 3.22979 | 0.264092 | 0.016 | 0.016 |

B) Minor glycan type

| species 1 | species 2 | F.Model | R2 | p.value | p.adjusted |
|---|---|---|---|---|---|
| Bmin | Dtren | 4.273389 | 0.2993955 | 0.016 | 0.016 |
| Bmin | Smic | 10.329241 | 0.5080977 | 0.003 | 0.003 |
| Bmin | Cgor | 4.999447 | 0.3571174 | 0.016 | 0.016 |
| Dtren | Smic | 9.618212 | 0.4902696 | 0.013 | 0.013 |
| Dtren | Cgor | 1.683928 | 0.1576132 | 0.203 | 0.203 |
| Smic | Cgor | 2.466919 | 0.2151335 | 0.112 | 0.112 |

C) Major glycan type (mannose, hybrid, complex)

| species 1 | species 2 | F.Model | R2 | p.value | p.adjusted |
|---|---|---|---|---|---|
| Bmin | Dtren | 7.312707 | 0.4223896 | 0.014 | 0.014 |
| Bmin | Smic | 24.810636 | 0.7127315 | 0.008 | 0.008 |
| Bmin | Cgor | 7.349677 | 0.4495304 | 0.021 | 0.021 |
| Dtren | Smic | 21.42005 | 0.6817319 | 0.004 | 0.004 |
| Dtren | Cgor | 2.645179 | 0.227148 | 0.137 | 0.137 |
| Smic | Cgor | 2.628443 | 0.2260357 | 0.139 | 0.139 |
